# Cerebral ischemia-induced genes are increased in acute schizophrenia: an opportunity for clinical translation of genomic research findings

**DOI:** 10.1101/158436

**Authors:** Hans W. Moises, Moritz Hess, Harald Binder

## Abstract

Schizophrenia is a brain disorder of unknown etiology. Brain imaging studies have revealed evidence for hypoperfusion of the frontal cortex (hypofrontality) and progressive brain volume reduction in schizophrenic patients. Mild cerebral ischemia (oligemia) has been postulated as a cause of the disorder. If the ischemia hypothesis for the adult brain is correct, genes induced by cerebral ischemia should be increased in the frontal cortex of schizophrenic patients during acute psychosis. Here, we show for the first time through a combined analysis of gene expression data from all the studies of the Stanley Brain Collection covering the Brodmann area 46 of the frontal cortex and employing the well-established Affymetrix HGU133a microarray platform that genes upregulated by cerebral ischemia are significantly overexpressed (4.5-fold) in the frontal cortex of acute schizophrenic patients (representation factor (RF) 4.5, *p* < 0.0002) and to a lesser degree in chronic patients (RF 3.9, *p* < 0.008) in comparison to normal controls. Neurodevelopmental-, repair-, inflammation- and synapse-related genes showed no significant change. The difference between acute and chronic schizophrenic patients regarding cerebral ischemia-induced genes was highly significant (RF 2.8, *p* < 0.00007). The results reported here are in line with evidence from biochemical, cellular, electroencephalographic, brain imaging, cerebral near-infrared spectroscopy, vascular, and genetic association studies. In summary, our genomic analysis revealed a clear ischemic signature in the frontal cortex of schizophrenia patients, confirming the prediction of the adult ischemia hypothesis for this disorder. This finding suggests new possibilities for the treatment and prevention of schizophrenia.

## INTRODUCTION

Schizophrenia is a severe mental disorder characterized by hallucinations, delusions, cognitive deficits, and a heterogeneous clinical course^1,2^. Brain imaging studies of this disorder have revealed a disturbance of brain function and a progressive decline in brain volume^3,4^. Schizophrenia has a strong genetic component with heritability estimates ranging from 64% to 90%.^5–7^. Environmental factors are involved in the onset and relapse.^7–9^

A large number of genes associated with schizophrenia have been implicated through genetic as well as genome-wide association studies.^10,11^ However, the field still faces the significant challenge of translating these findings into novel understandings of the pathophysiology of schizophrenia and possible drug targets.^12,13^ Several hypotheses have been put forward^13–22^. Recently, Moises et al. (2015) proposed a vascular-ischemia hypothesis for schizophrenia of the adult brain based on functional-genomic analyses of susceptibility genes for the disorder^22^.

If the adult ischemia hypothesis is correct, genes induced by cerebral ischemia should be significantly higher expressed in the postmortem brains of acute schizophrenic patients compared to normal controls or to chronic patients. The aim of the present study was to test this prediction. Here, by combining the gene expression microarray data from two genome-wide expression studies of postmortem samples from the Stanley Online Genomics Database covering the frontal cortex (Brodmann area 46), we find that cerebral ischemia-induced genes are indeed significantly overexpressed in the frontal cortex of acute schizophrenic patients.

## MATERIAL AND METHODS

### Postmortem material and gene expression data

To investigate the association of acute schizophrenia and frontal cerebral gene expression, we combined expression data from two studies obtained from the Online Genomics Database of the Stanley Medical Research Institute (SMRI) [https://www.stanleygenomics.org/]. Details of the original microarray gene expression studies are shown in Supplementary Table S1.

The RMA normalized expression data were downloaded from the Online Genomics Database of the SMRI. Only studies investigating gene expression in the Brodmann area 46 (BA46) were considered. We selected the data sets "AltarC" and Kato" since they covered all donors from the Stanley Brain Collection (A and C), investigated with the popular Affymetrix HGU133a microarray platform (for details, see supplementary Table S1). Acute schizophrenia was defined as having a psychotic feature and definite or probable exacerbation. Chronic schizophrenia was defined as probably not having an exacerbation.

### Functional gene sets

Functional gene sets induced by cerebral ischemia, neurodevelopment, post-ischemiaic repair, and synapse-related genes were obtained by extensive literature mining as described elsewhere^22^. Additionally, an inflammation-related gene set of 500 genes was used in the present study. It consisted of 36 inflammation-related genes extracted from the Entrez Genes database of homo sapiens (HS) using the search term *‘inflammation* OR *inflammatory*’^23^, 179 genes induced by lupus erythematosus^24–30^, 266 genes induced by autoimmunity or multiple sclerosis^26,31^, and 252 genes from the chemokine signaling and lupus erythematosus pathways of the KEGG database^32^. Removal of duplicates resulted in 500 inflammation-related genes.

## METHODS

### Matching of patients and controls

Patients and controls were matched by cluster analysis based on Euclidean distance. Confounding variables were considered in the following order: brain collection (A, C), sample preparation (frozen, fixed), and brain pH. Factors were weighted based on the expected influence on gene expression. Clustering results were compared with the optimal matching approach implemented in MatchIt (method = ‘optimal’).^33^ In particular, we compared the average intra-cluster Euclidean distance in matching pairs generated by our approach and matching pairs generated by MatchIt. Since our approach resulted in a lower distance, we used the respective cluster in the subsequent analysis. The clustering does generally show good agreement in terms of cohorts, sample preparation, brain pH, and age at death (Supplementary Figure 1 (Clustering diagram) and Supplementary Table S2 (Clustering Table). Matching was conducted using the statistical programming language R v.3.3.0.^34^

**Figure 1.**
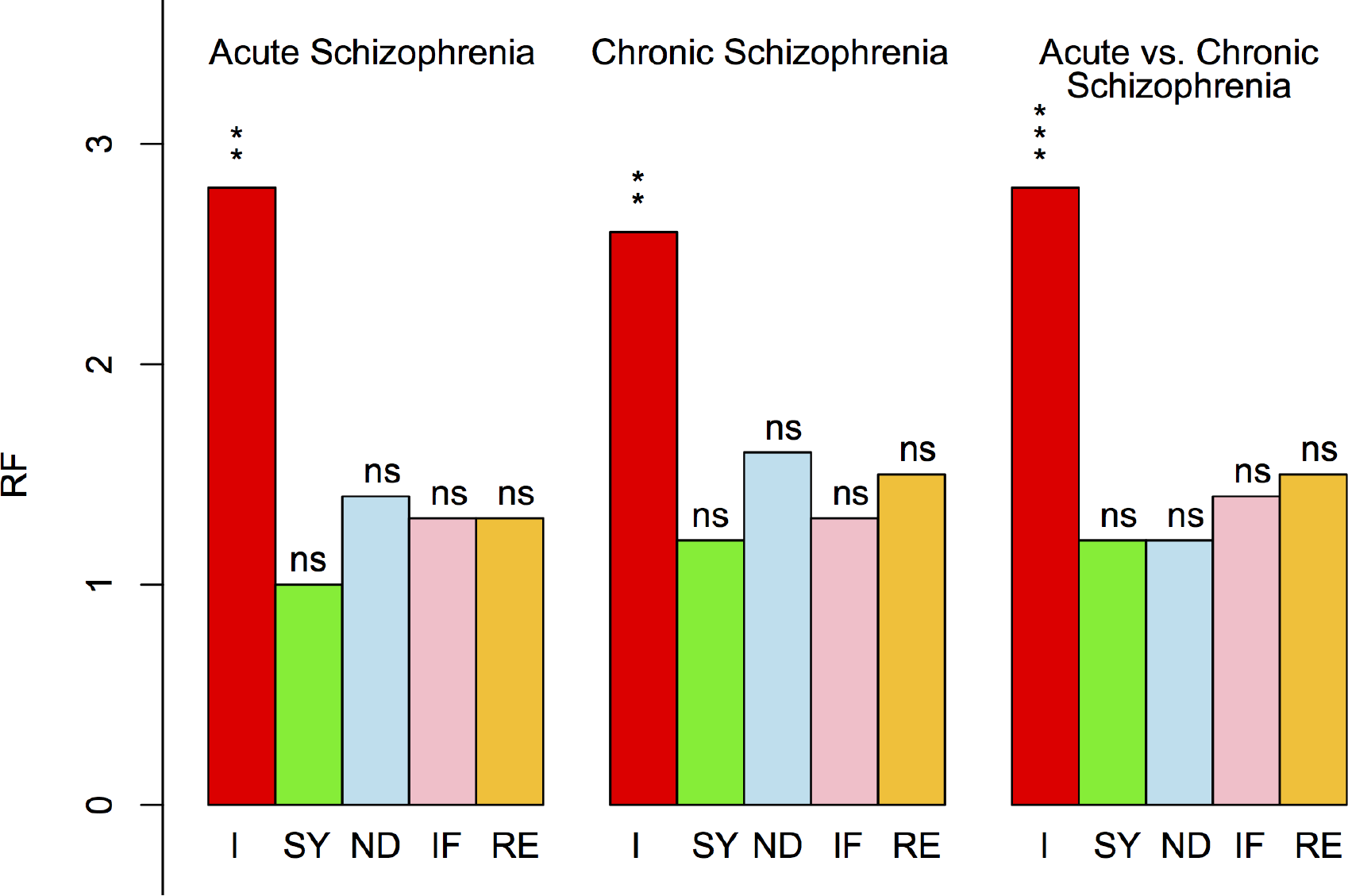
Functions of the top 100 differentially expressed genes in the postmortem brains of adult patients suffering from acute or chronic schizophrenia. I = genes differentially expressed due to cerebral ischemia, SY = synapse-related genes, ND = genes differentially expressed during neurodevelopment, IF = inflammation-induced genes, RE = repair genes (i.e., genes involved in neuronal repair). Level of significance: * < 0.01, ** < 0.001, *** < 0.0001.

### Normalization of expression data and analysis of differential gene expression

The odds of being diagnosed with acute schizophrenia (AS) compared to (i) being diagnosed with chronic schizophrenia (CS) or (ii) not being diagnosed with schizophrenia (control), dependent on expression levels of a single gene, were investigated using univariate conditional logistic regression. The RMA normalized expression data measured on the Affymetrix HGU133a platform were employed. Expression data were aggregated on gene levels, leading to expression data of 13 516 genes that were investigated. Strata in the conditional logistic regression models corresponded to the matching groups (Supplementary Table S2). Significance of odds ratios between two groups was calculated via a likelihood-ratio test. In total, three group comparisons were conducted: AS vs. control, CS vs. control, and AS vs. CS. No adjustment for multiple testing was performed. The results are provided in Supplementary Table S3. The top 100 differentially expressed genes across all three comparisons were inspected in detail. Genes were ranked by the combined *p*-values from all three group comparisons and only the top 100 differentially expressed genes were retained (Supplementary Figure S2). The *p*-values were combined using Fisher’s method. All statistical analyses were conducted using the statistical programming language R v. 3.3.0.^34^

### Functional genomic analysis

The number of overlaps between the top 100 differentially expressed genes and functional gene sets was determined by the intersect function implemented in R.^34^ Genes were considered differentially expressed in the respective comparison (e.g., AS vs. control) if they showed at least a slight indication of differential expression (no white color in Supplementary Figure S2). Probes without gene IDs were discarded. Statistical significance and confidence intervals for the number of matches between the 100 differentially expressed genes and each functional gene set were determined using separate Fisher’s exact tests (one sided) as implemented in R.^34^ The validity of this statistical approach was verified using a custom genome resampling test (described in detail elsewhere^22^), which gave the same results. The family-wise error rate (FWER) was controlled at the 1% level using Bonferroni’s correction.

## RESULTS

The heatmap of the top 100 differentially expressed genes from the postmortem brains of adult schizophrenic patients is displayed in Supplementary Figure S2. The figure shows, for example, that NOS1AP (row 15), a gene involved in the synthesis and binding of nitric oxide and vasodilatation, is downregulated in the acute compared to chronic schizophrenic brain. The same holds true for MAOB (row 16), a gene encoding monoamine oxidase type B, which is responsible for the degradation of dopamine, a vasoconstrictive neurotransmitter.^22^

Among the top 100 genes, 63 genes were differentially expressed in acute schizophrenic patients (37 upregulated and 26 downregulated) and 95 genes in chronic schizophrenics (27 upregulated and 68 downregulated) compared to normal controls. Furthermore, 90 of the top 100 genes were differentially expressed (62 upregulated and 28 downregulated) in acute compared to chronic schizophrenic patients.

### Functional genomic analysis

The results of the functional genomic analysis of the top 100 differentially expressed genes are provided in Tables 1–3 and depicted in Figure 1. The results show that genes induced by cerebral ischemia are significantly overexpressed in the brains of patients suffering from acute or chronic schizophrenia. Furthermore, the comparison between acute and chronic schizophrenic patients revealed a highly significant difference, specifically for genes induced by cerebral ischemia.

### Analysis of subgroups

Subgroup analyses of ischemia genes (i.e., testing genes that are upregulated or downregulated by cerebral ischemia for intersection with upregulated or downregulated genes in the brains of schizophrenic patients) revealed significant results only for genes upregulated by ischemia. Compared to the controls, acute schizophrenic patients showed 4.5 times higher expression of brain ischemia-induced genes (representation factor (RF) of 4.5; *p* = 0.0002), while for chronic patients the expression was 3.9 times higher (RF of 3.9; *p* = 0.008). Moreover, patients with acute schizophrenic psychoses differed significantly from chronic patients regarding genes upregulated by cerebral ischemia (RF of 3.7, *p* = 0.0001). By contrast, the results for down-regulated genes were non-significant, with an RF of 1.7 (*p* = 0.45), 1.9 (*p* = 0.19), and 1.6 (*p* = 0.47) for acute, chronic, and acute vs. chronic schizophrenia, respectively.

### Dose-response relationship

The data show a dose-response relationship between the expression of genes upregulated by cerebral ischemia and the disease status of schizophrenic patients (Figure 2).

**Figure 2.**
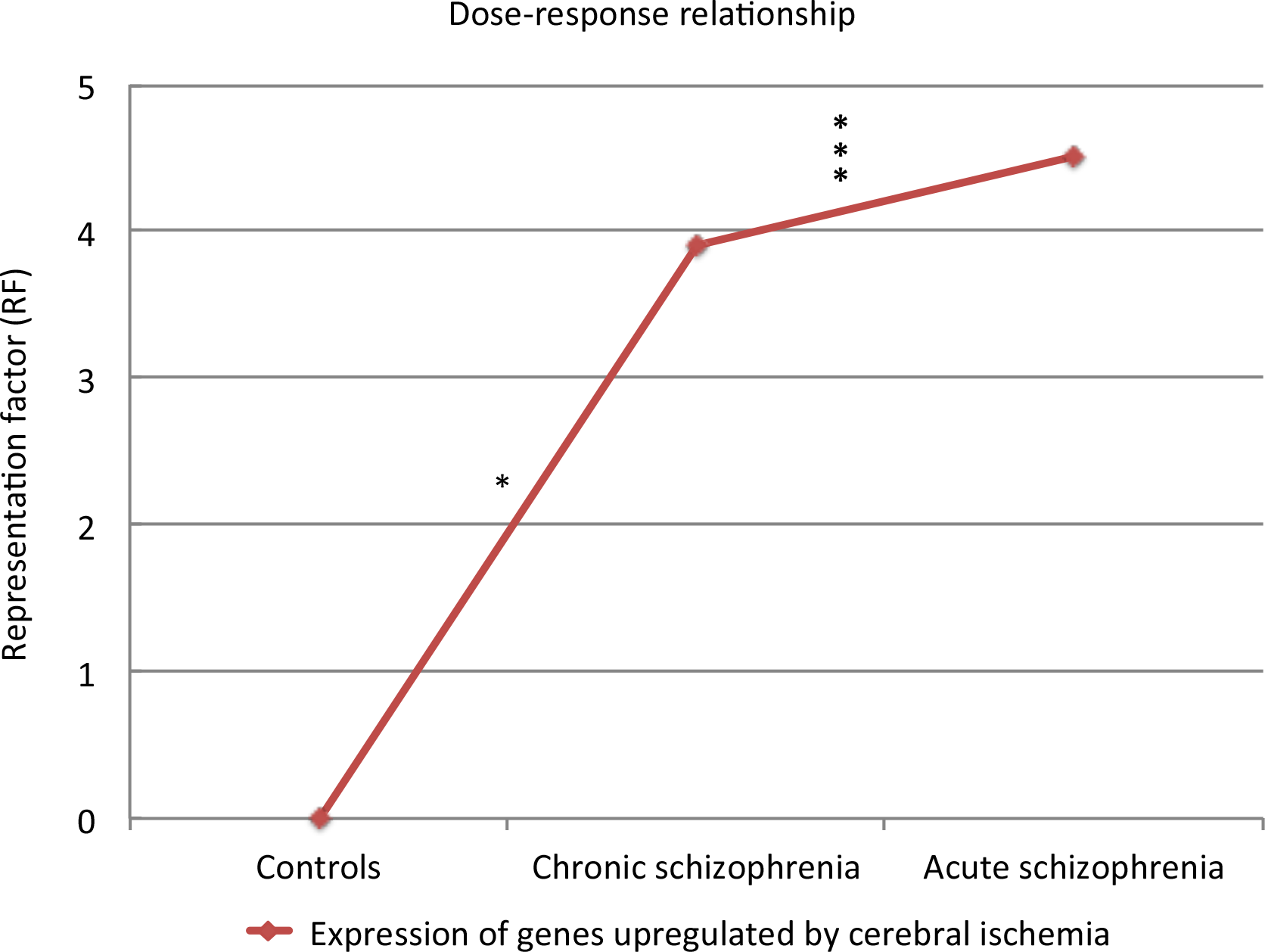
Dose-response relationship between expression of genes induced by cerebral ischemia and disease status in schizophrenic patients. RF = representation factor compared to normal controls. Level of significance: * ≤ 0.01, *** ≤ 0.0001.

## DISCUSSION

The starting point of the present investigation was the prediction that, if the adult ischemia hypothesis of schizophrenia^22^ were correct, evidence for cerebral ischemia should be detectable in the brains of acute schizophrenic patients compared to normal controls as well as to chronic schizophrenic patients. To test this prediction, genome-wide expression data from two postmortem studies available from the Genomics Database of the SMRI were pooled, and the differential expression was analyzed separately for acute and chronic schizophrenic patients. The analysis of the overlap between the top 100 differentially expressed genes (heatmap, Supplementary Figure S2) and functional gene sets gave a surprisingly clear result. Genes induced by cerebral ischemia were significantly overexpressed in the brains of acutely ill schizophrenic patients compared to the matched controls (Figure 1 and Table 1). The same held true for chronic schizophrenic patients, although to a lesser extent as indicated by the lower RF of 2.6, compared to 2.8 in acute patients (Tables 1 and 2; Figure 2) and an RF of 3.9 compared to 4.5 for genes upregulated by cerebral ischemia. In contrast, gene sets involved in synaptic functioning, brain growth (neurodevelopment), inflammation, and repair showed no significant differences from the controls (Figure 1).

**Table 1.** Intersection analysis between functional gene sets and the top 100 genes differentially expressed in the postmortem brains of acute schizophrenic patients versus normal controls (AS.Control).

| Functions | *N* | SZ<br>*N* | E<br>(N) | O<br>(N) | E<br>(%) | O<br>(%) | RF | CI | Nominal<br>*p* ≤ | Bonferroni<br>corrected *p* ≤ |  |
| --- | --- | --- | --- | --- | --- | --- | --- | --- | --- | --- | --- |
| I | 1673 | 63 | 5 | 14 | 8 | 22.2 | 2.8 | 9-63 | 0.0003 | 0.001 | ** |
| SY | 1977 | 63 | 6 | 6 | 9.5 | 9.5 | 1.0 | 3-63 | 0.53 | 1.00 | n.s. |
| ND | 3211 | 63 | 10 | 13 | 15.3 | 20.6 | 1.4 | 8-63 | 0.14 | 0.70 | n.s. |
| IFL | 500 | 63 | 1 | 2 | 2.4 | 3.2 | 1.3 | 0-63 | 0.44 | 1.00 | n.s. |
| RE | 3319 | 63 | 10 | 13 | 15.8 | 20.6 | 1.3 | 8-63 | 0.17 | 0.85 | n.s. |

I = genes differentially expressed by cerebral ischemia, SY = synapse-related genes, ND = genes differentially expressed during neurodevelopment, IFL = inflammation-induced genes, RE = repair, genes involved in neuronal repair, RF = representation factor. Level of significance: \* ≤ 0.01, \*\* ≤ 0.001, \*\*\* ≤ 0.0001.

**Table 2.**
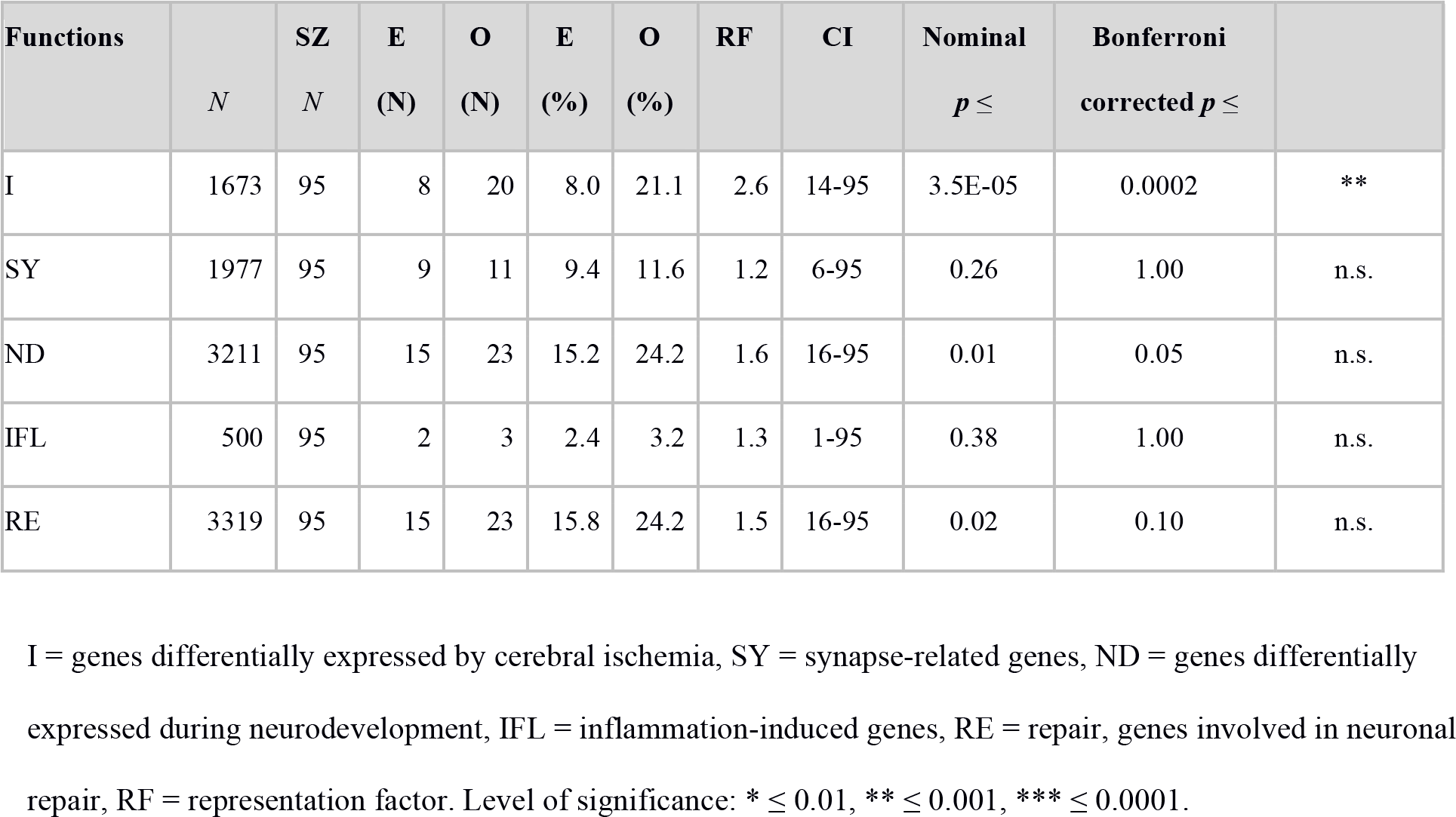
Intersection analysis between functional gene sets and the top 100 genes differentially expressed in the postmortem brains of chronic schizophrenic patients versus normal controls (CS.Control).

| Functions | *N* | SZ<br>*N* | E<br>(N) | O<br>(N) | E<br>(%) | O<br>(%) | RF | CI | Nominal<br>*p* ≤ | Bonferroni<br>corrected *p* ≤ |  |
| --- | --- | --- | --- | --- | --- | --- | --- | --- | --- | --- | --- |
| I | 1673 | 95 | 8 | 20 | 8.0 | 21.1 | 2.6 | 14-95 | 3.5E-05 | 0.0002 | ** |
| SY | 1977 | 95 | 9 | 11 | 9.4 | 11.6 | 1.2 | 6-95 | 0.26 | 1.00 | n.s. |
| ND | 3211 | 95 | 15 | 23 | 15.2 | 24.2 | 1.6 | 16-95 | 0.01 | 0.05 | n.s. |
| IFL | 500 | 95 | 2 | 3 | 2.4 | 3.2 | 1.3 | 1-95 | 0.38 | 1.00 | n.s. |
| RE | 3319 | 95 | 15 | 23 | 15.8 | 24.2 | 1.5 | 16-95 | 0.02 | 0.10 | n.s. |

I = genes differentially expressed by cerebral ischemia, SY = synapse-related genes, ND = genes differentially expressed during neurodevelopment, IFL = inflammation-induced genes, RE = repair, genes involved in neuronal repair, RF = representation factor. Level of significance: \* ≤ 0.01, \*\* ≤ 0.001, \*\*\* ≤ 0.0001.

**Table 3.** Intersection analysis between functional gene sets and the top 100 genes differentially expressed in the postmortem brains of acute versus chronic schizophrenic patients (AS.CS).

| Functions | *N* | SZ<br>*N* | E<br>(N) | O<br>(N) | E<br>(%) | O<br>(%) | RF | CI | Nominal<br>*p* ≤ | Bonferroni<br>corrected *p* ≤ |  |
| --- | --- | --- | --- | --- | --- | --- | --- | --- | --- | --- | --- |
| I | 1673 | 90 | 7 | 20 | 8.0 | 22.2 | 2.8 | 14-90 | 1.5E-05 | 0.00007 | *** |
| SY | 1977 | 90 | 8 | 10 | 9.4 | 11.1 | 1.2 | 6-90 | 0.31 | 1.00 | n.s. |
| ND | 3211 | 90 | 14 | 22 | 15.3 | 24.4 | 1.6 | 15-90 | 0.01 | 0.05 | n.s. |
| IFL | 500 | 90 | 2 | 3 | 2.4 | 3.3 | 1.4 | 1-90 | 0.35 | 1.00 | n.s. |
| RE | 3319 | 90 | 14 | 22 | 15.8 | 24.4 | 1.5 | 15-90 | 0.02 | 0.10 | n.s. |

I = genes differentially expressed by cerebral ischemia, SY = synapse-related genes, ND = genes differentially expressed during neurodevelopment, IFL = inflammation-induced genes, RE = repair, genes involved in neuronal repair, RF = representation factor. Level of significance: \* ≤ 0.01, \*\* ≤ 0.001, \*\*\* ≤ 0.0001.

Importantly, the results of the combined genome-wide expression analysis presented in this study are consistent with the occurrence of functionally relevant hypoperfusion of the frontal cortex during acute schizophrenic psychosis (meta-analyses^35,36^). Furthermore, they agree with the recent findings by Katsel et al. (2017) that genes involved in angiogenesis-related pathways are downregulated in brain samples from schizophrenic patients, suggesting an impaired cerebrovascular plexus^37^. In line with these findings, schizophrenic patients and their healthy family members show an increase of the anti-angiogenic factor "soluble fms-like tyrosine kinase-1 (sFlt-1)" in their plasma.^38,39^

Our findings are not at variance with the results from the postmortem expression studies performed by other researchers. Prabakaran et al. (2004) performed an impressive NMR-based metabolomic study and reported upregulation of pathways associated with oxygen and reactive oxygen species metabolism, significant downregulation of pathways involved in oxidative phosphorylation and energy pathways, and significantly raised lactate levels in schizophrenic brains. These authors emphasized mitochondrial dysfunction as a possible cause of schizophrenia, without ruling out hypoxic events and oxidative stress.^40^ The investigation by Altar et al. (2005) revealed downregulation of genes encoded for mitochondrial energy metabolism^41^, Iwamoto et al. (2005) reported global down-regulation of mitochondrial genes.^42^ Hakak et al. (2001) discovered differential expression of myelination-related genes in schizophrenia suggesting a disruption of oligodendrocyte function^43^, a finding that has been confirmed by other expression and proteomic studies^44^. The myelin-related findings in schizophrenia might be explained by the selective vulnerability of oligodendrocytes to ischemia.^45^

Mistry et al. (2013) performed a comprehensive meta-analysis of seven expression studies from the frontal cortex of schizophrenic patients. They obtained a signature of 39 upregulated and 86 downregulated genes that was consistent with alterations in energy metabolism and synaptic transmission.^46^

Moises et al. (2015)^22^ employed the same gene sets as in the present study to conduct a functional-genomic analysis of the results obtained by Mistry et al.^46^, and reported an upregulation of vascular-ischemic genes and a downregulation of synaptic as well as neurodevelopmental/repair genes.^22^ Although the results of Moises et al.’s (2015) genomic postmortem study are in agreement with those of the present investigation, the latter are more clear-cut than the former, possibly due to being based on the original expression data instead of the small number of replicated genes identified by Mistry et al. in their meta-analysis.^46^ Additional differences include the matching procedure and criteria, two instead of seven studies being investigated, and the differentiation between acute and chronic schizophrenia.

Taken together, nearly all postmortem studies have so far either obtained direct^22,40^ or indirect^41–44^ evidence consistent with mild cerebral ischemia in the adult brain of schizophrenic patients. Most importantly, the results presented here are in agreement with established findings from five different fields of schizophrenia research: (1) biochemical, cellular, electroencephalographic, and clinical signs of cerebral hypoxia/ischemia^47–50^, (2) oxygen insufficiency of the frontal cortex in acute schizophrenic patients detected by near-infrared spectroscopy (NIRS)^51,52^, (3) hypofrontality observed by brain imaging^36^, (4) vascular studies (e.g.,^53,54^), and (5) genetic association studies.^22^ For example, Reif et al. were able to demonstrate that NOS1, a risk gene for schizophrenia, is involved in reducing brain oxygenation in patients.^55^ Further references and information is given in the supplementary section.

To explain hypofrontality in the brains of schizophrenic patients, three categories of hypotheses have been put forward: (i) neurodevelopmentally caused brain lesions, (ii) frontal hypoactivation, and (iii) hypoperfusion.

The neurodevelopmental brain lesion theory postulates that cerebral hypoperfusion is not the cause of the disease but a consequence of a yet undiscovered brain lesion. It is well known that cerebral metabolism and cerebral blood flow (CBF) are tightly coupled. Hence, reduced CBF might simply reflect a subtle neurodevelopmental brain lesion (see Weinberger^56,57^ and Schmidt-Kastner^18,58^). The hypoactivation hypothesis resembles the lesion theory but replaces the postulated neurodevelopmental brain lesion with schizophrenic psychopathology. In other words, schizophrenic patients perform cognitive tasks more poorly than the controls and hence activate their frontal cortex to a lesser degree, resulting in a reduced need for oxygenation and CBF (see Frith^59^ and Bullmore^60^). By contrast, hypoperfusion hypotheses see the reduced CBF as causing the reported neuronal-glial dysfunction and the observed psychopathological symptoms in schizophrenic patients (see, e.g., Bachneff’s local circuit neurons hypothesis^61^, Hanson and Gottesman’s genetic-inflammatory-vascular hypothesis^14^, Lopes et al.’s angiogenesis^62^, Katsel et al.’s impaired cerebrovascular plexus^37^, and Moises et al.’s adult vascular-ischemia hypothesis.^22^

It is difficult to see how brain lesions caused during neurodevelopment or how functional hypoactivation of the frontal cortex during psychosis could increase the expression of genes induced by cerebral ischemia in the adult brain or produce the other signs of hypoxia/ischemia in schizophrenic patients listed above. The hypoperfusion hypothesis is consistent with the results of a recent brain imaging study in schizophrenic patients showing in the left lateral prefrontal cortex (LPFC) and the right anterior cingulate cortex (ACC) significantly reduced CBF despite normal grey matter volume (GMV).^63^ The authors concluded that such changes in CBF are unlikely to be caused by GMV alterations.^63^ Moreover, the neurodevelopmental lesion and the hypoactivation hypotheses are not able to explain the progressive brain volume reduction seen in schizophrenic patients (meta-analysis^64^).

In summary, our finding of a significant increase in the expression of ischemia-induced genes in the brains of acute and – to a lesser extent – of chronic schizophrenic patients confirms the prediction of the adult vascular-ischemia hypothesis.^22^ Furthermore, this hypothesis is able to explain many diverse biochemical, cellular, electroencephalographic, and clinical findings, and the results of cerebral NIRS, brain imaging, vascular and genetic association studies^22^ (see the supplementary discussion).

We conclude that acute schizophrenia appears to be accompanied by mild cerebral ischemia, suggesting that the latter might be the cause of the disorder, or at least play a role in its pathogenesis. Hence, in acute schizophrenic patients, supply of glucose^65^ or oxygen ^66^, vasodilatation (e.g., by nicotine agonists^67^ or by omega-3^68,69^), and improvement of CBF (e.g., by erythropoietin ^70,71^, ginkgo ^72,73^ or nimodipine ^74,75^) should have a positive effect on the symptomatology and outcome of the disease.

## ACKNOWLEDGEMENTS

We are most grateful to the Stanley Online Genomics Database (https://www.stanleygenomics.org) for providing the expression data obtained from collaborators who investigated samples from the Brain Bank of the Stanley Medical Research Institute (SMRI). The collaborators are Tadafumi Kato (Japan) and C. Anthony Altar (USA). Without their data and the SMRI, our analysis and this publication would not have been possible.

